# A litmus test for harmonized mosquito monitoring across Europe and Africa

**DOI:** 10.1101/2020.01.30.927020

**Authors:** Ignazio Graziosi, Carles Aranda, Fabrizio Balestrino, Romeo Bellini, Núria Busquets, Mammadou Coulibaly, Andrea Crisanti, Diawo Diallo, Mawlouth Diallo, Alioune Gaye, Moussa Guelbeogo, Aleksandra Ignjatović-Ćupina, Sebastián Napp, Sagnon N’Falé, Dušan Petrić, Paola Pollegioni, Alekos Simoni, Marija Zgomba, Ruth Müller

## Abstract

The accelerating rate of outbreaks from mosquito borne diseases are urging the development of updated and effective tools for the surveillance of insect populations and their larval habitats. Harmonized field protocols help to build a comprehensive picture on species-specific vector ecology and generate important knowledge for implementing coordinated mosquito surveillance programs at regional scales and across continents. In this study, we test the efficiency and potential barriers of available harmonized protocols from earlier EU project VectorNet. As a kind of litmus test for such protocols, we specifically aim to capture the ecoregional variation of breeding site characteristics and population density of five mosquito vectors in Europe and Africa. As expected, the five species considered show different aquatic habitat preferences. *Culex pipiens, Aedes albopictus* in Europe and *Ae. aegypti* in Africa select breeding habitats within specific volume classes, while *Anopheles gambiae* and *An. coluzzii* may select breeding habitats based on seasonal availability. Population densities in aquatic habitats greatly varied across species and countries, but larval production sites of *Ae. albopictus* generate populations with higher ratio of pupae compared to the other species. This result underlines the fundamental ecological difference between the selected vector species disregarding of the ecoregion. Mean water temperatures had limited variation across species and higher among countries. Understanding the ecology of native and non-native mosquito vectors is key in evaluating transmission risks for diseases such as West Nile, chikungunya and dengue fevers, zika and malaria. The available harmonized field protocols are a valuable tool for achieving homogeneous mosquito surveillance in Europe and Africa.

## Introduction

The global trend of mosquito-borne diseases is dramatically rising, with recrudescence of some well-known pathogens and accelerating outbreaks of emerging diseases in multiple regions (Jones et al. 2008). mosquito-borne diseases are a major burden to human health, thus determining pervasive health and socio-economic costs to local communities (Gubler 1998). We are witnessing the rising of once neglected arboviral tropical diseases carried by native and invasive mosquitoes, namely dengue fever, Zika, chikungunya and West Nile, which are quickly becoming pandemic and pose a current threat for Europe and Africa (Benelli and Mehlhorne 2016, Weaver et al. 2018). Accelerated human mobility facilitates the movement of infected individuals (Sarma et al. 2018), global trade is intensifying introduction of non-native vectors, and changing climates and anthropization are creating suitable habitats for mosquitoes, thus magnifying proliferation and risk of transmission (Derraik and Slaney 2007). Furthermore, malaria incidence is locally increasing in high-risk areas of Africa and South America and resurging unexpectedly in Greece (Danis et al. 2011, Whitty and Ansah, 2019).

As a result of anthropogenic-driven mosquito proliferation and climatic adaptations we may expect the recrudescence of old and new infectious diseases vectored by several species of mosquitoes (Diptera: Culicidae) world-wide (Hotez 2016), including *Culex* and *Aedes* (Culicinae) and *Anopheles* (Anophelinae). *Culex pipiens* Linnaeus is a common mosquito with world-wide distribution, a competent vector for West Nile and Japanese encephalitis viruses which contributed to West Nile outbreaks in Southern Europe and North America (Wispelaere 2017, Marini et al. 2020). The highly invasive *Aedes albopictus* (Skuse) is expanding its non-native range in Europe thanks to exceptional ecological plasticity, and is increasingly alarming health organizations and surveillance networks for its ability to cause dengue, Zika and chikungunya outbreaks (Aranda et al. 2018, Giron et al. 2019, Javelle et al. 2019). Congeneric *Ae. aegypti*, the vector of yellow fever virus in Africa, is also competent for dengue, chikungunya and Zika viruses, thus magnifying infection risks in Europe, where the species was also detected (Akiner et al. 2016). *Anopheles gambiae* Giles and *An. coluzzii* Coetzee & Wilkerson are the most widespread species causing malaria in Sub Saharan Africa, with the former preferring habitats in the African humid tropics and the latter dryer habitats, typically in the Sahel (Lanzaro and Lee 2013, Coetzee et al. 2013).

Understanding the ecology of native and non-native mosquito vectors, and how habitat availability define their distribution, is key in evaluating transmission risks. Environmental parameters of mosquito breeding site (MBS) shape insect development and ultimately affect their vector competence. The productivity of MBS is determined by water temperature, volume of breeding environment, predator density or presence of organic matter (Bevins 2008, Okech et al. 2007, Schneider et al. 2011). The variation of the productivity of MBS within ecoregions is still unclear (Pascoe et al. 2019); though the characterization of such variations would add fundamental knowledge on population dynamics and structures (Okogun et al. 2003, Vallorani et al. 2015).

The success of managing emerging and resurging mosquito-borne diseases relies on designing effective integrated vector management (IVM) tools, which include active vector monitoring (Fernandes et al. 2018). The effectiveness and reliability of pan-regional systems for vector monitoring requires the development of homogeneous methodologies. Recent EU research networks (Vector Net, AIM Cost) aimed to standardize procedures for the surveillance of arthropod vectors in Europe in order to provide standardized datasets (Versteirt et al. 2017, Medlock et al. 2018). Suggestions on the use of harmonised methods, equipment and field protocols for sampling have been published (ECDC and EFSA 2018). However, a litmus test of such protocols is needed to evaluate the realistic potential of methodological standardization and challenges in different ecoregional settings (Vernick 2017).

Within the framework of the EU funded InfraVec2 consortium, we tested the efficiency and potential barriers of available harmonized protocols from earlier EU project VectorNet. As a kind of litmus test for such protocols, we specifically aimed to capture the ecoregional variation of MBS and population densities of five target mosquito vectors in Europe and Africa.

## Material and methods

We conducted a 6-country study describing the variation in natural vector larval populations to test harmonized field protocols and assessed regional variation of MBS parameters for five mosquito species: (1) *Cx. pipiens* in Spain, Italy and Serbia, (2) *Ae. albopictus* in Spain and Italy, (3) *Ae. aegypti* in Senegal, (4) *An. gambiae* in Senegal and Burkina Faso, and (5) *An. coluzzii* in Mali.

At first, we jointly developed standard operating procedures (SOP) for a MBS survey based on the VectorNet field protocols and guidelines from the European Centre for Disease Prevention and Control (ECDC and EFSA 2018). The agreed SOP was tested in all six countries: in Catalonia, Spain, in the Emilia Romagna region of Italy, in Novisad and Voivodina area in Serbia, Dakar and Kedougou province in Senegal, Ouagadougou and Banfora areas in Burkina Faso, and Bamako and Kati districts in Mali (Fig. 1). Potential MBS were surveyed in 2018 and 2019 during mosquito peak season (July to early August in Europe, late September to early November in Africa) except from Mali, which was surveyed at the end of the peak season (late December).

**Fig. 1.**
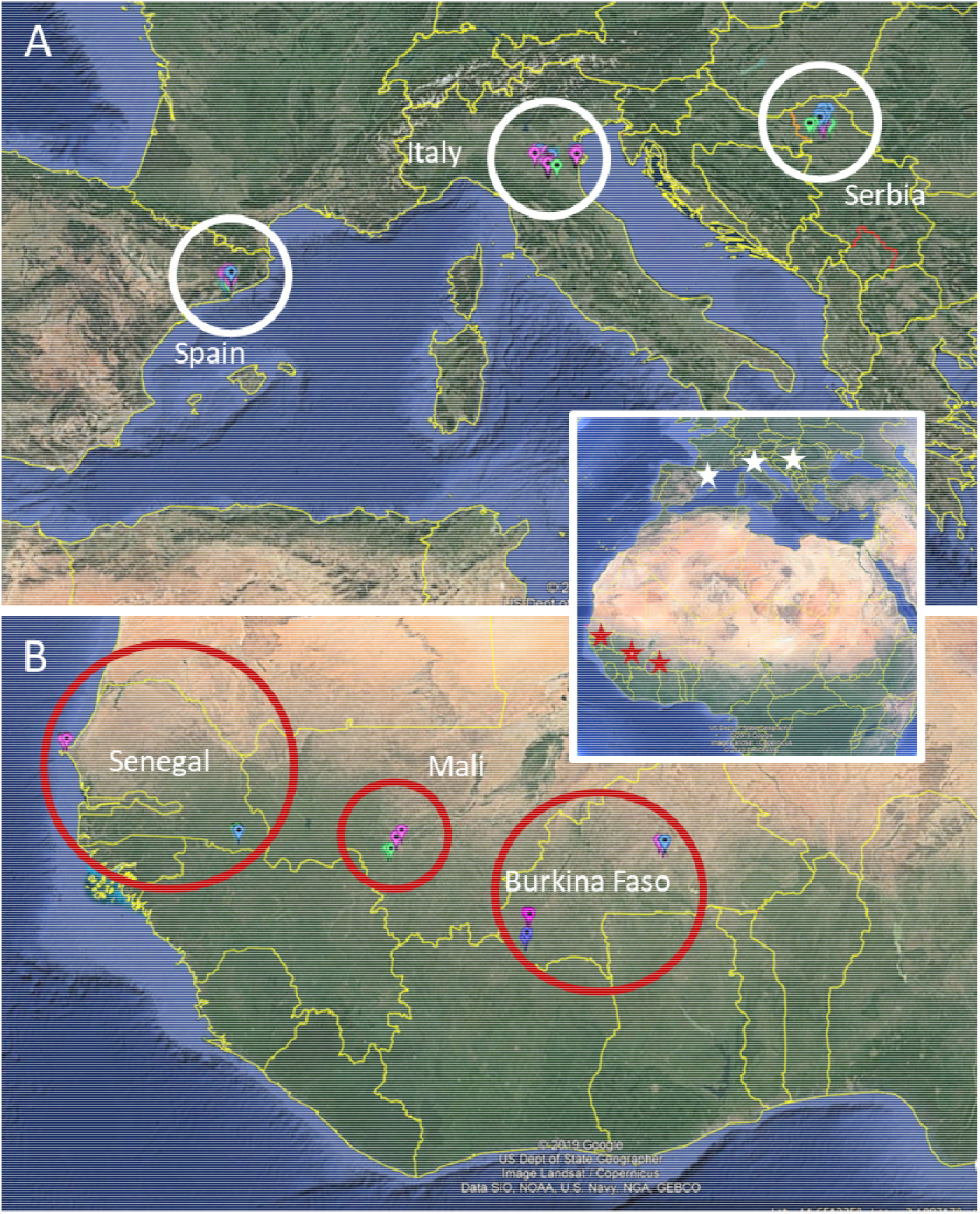
Harmonized mosquito monitoring protocols were tested in six countries located in: A) Europe (Spain, Italy, Serbia) and B) Sub-Saharan Africa (Senegal, Burkina Faso and Mali). (Imagery: Google Earth). Locations of mosquito breeding sites are available for download at: https://infravec2.eu/mosquito-survey-data/

In order to systematically comprise MBS variation across different landscapes, we surveyed potential MBS in urban (city), sub-urban (suburbs), rural (cropland) and natural habitats (rivers, wetlands, woodlands). For each MBS (mosquito-positive site), geographic coordinates were collected using hand-held GPS devices when available, or smartphones, georeferenced and displayed using Google Earth, and data from the field harmonized using MS Excel. The volume of MBS was assessed by measuring or by estimating the approximate dimensions of the water container/body of water. MBS were assigned to 6 volume classes: 1 (<0.5 L), 2 (0.5-5.0 L), 3 (5-50 L), 4 (50-200 L), 5 (200-2000 L), 6 (>2000 L). The type of MBS was also noted (i.e. river, pond, puddle, pot, tire, or other). We measured water temperature in the tray immediately to avoid heating using hand-held electronic thermometers with accuracy between 0.1-0.5°C.

In addition, we determined the mosquito population density and population structure in each MBS by using a dipper (or pipette for smaller breeding sites) to transfer 1L of water from the site into a plastic tray (or all the water for breeding sites <1L). We counted the number of young larvae (L1-L2), mature larvae (L3-4), and pupae (P) in the tray. Larvae and pupae were identified to species based on macro-morphology and habit. We also calculated the total number of mosquitoes T=L1-2+L3-4+P, and a pupal ratio R= P/T. For larger MBS, the water sampling and counting of larvae and pupae were replicated ten consecutive times. Means, standard deviations (STD), coefficient of variation (CV) and standard errors (SE) were calculated to assess variations of population density, population structure and MBS temperature.

## Results

In total, 191 MBS were assessed (Table 1): *Cx. pipiens* was collected from a total of 69 MBS (3 countries, Europe), *Ae. albopictus* from 35 MBS (2 countries, Europe), *Ae. aegypti* from 19 MBS (1 country, Africa), *An. gambiae* from 50 MBS (2 countries, Africa) and *An. coluzzii* from 18 MBS (1 country, Africa). Exact locations of MBS are available for download at: https://infravec2.eu/mosquito-survey-data/. We detected species-specific preferences for certain physical shape and volume of MBS: *Cx. pipiens* was preferentially found in canals, manholes, and streams, *Ae. albopictus* in flower pots or barrels used for common garden irrigation, *Ae. aegypti* in tires, *An. gambiae* in clay quarries and puddles, and *An. coluzzii* in puddles near river beds. The volume of MBS ranges from class 2 to 6 for *Cx. pipiens*, 1 to 4 for *Ae. albopictus*, 1 to 3 for *Ae. aegypti*, 3 to 5 for *An. gambiae*, and only class 3 for *An. coluzzii*. The characteristics and volume classes of MBS preferred by *Cx. pipiens*, *Ae. albopictus* and *An. gambiae* slightly vary among countries (Table 2, Fig. 2). The lowest temperature values were found in MBS of *Cx. pipiens*, and the highest in MBS of two Aedes species. The overall temperature in MBS varies only slightly across surveyed countries in Europe and Africa, ranging between 26.3 to 27.6 °C with an overall STD of 3.3 and CV of 12.5% (Table 3). The only exception is the mean temperature of the MBS of *An. gambiae*, which differs strongly between countries: in Burkina Faso the mean MBS temperature is 2°C cooler if compared to values of the respective MBS in Senegal (26.1°C and 28.1°C respectively, Fig. 3).

**Table 1.**
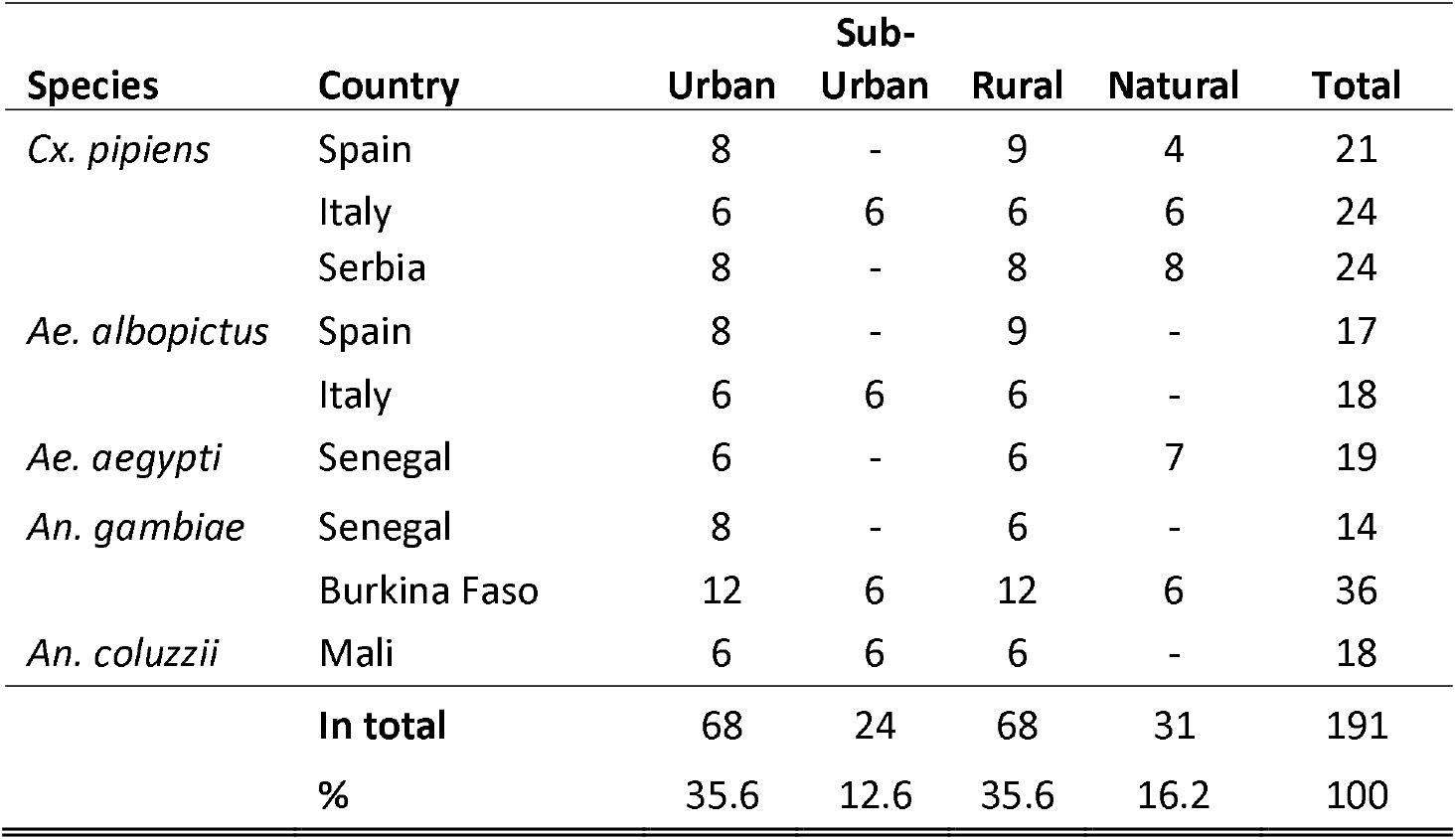
Mosquito breeding sites within a six-country survey conducted in Europe and Africa across different landscapes.

**Table 2.**
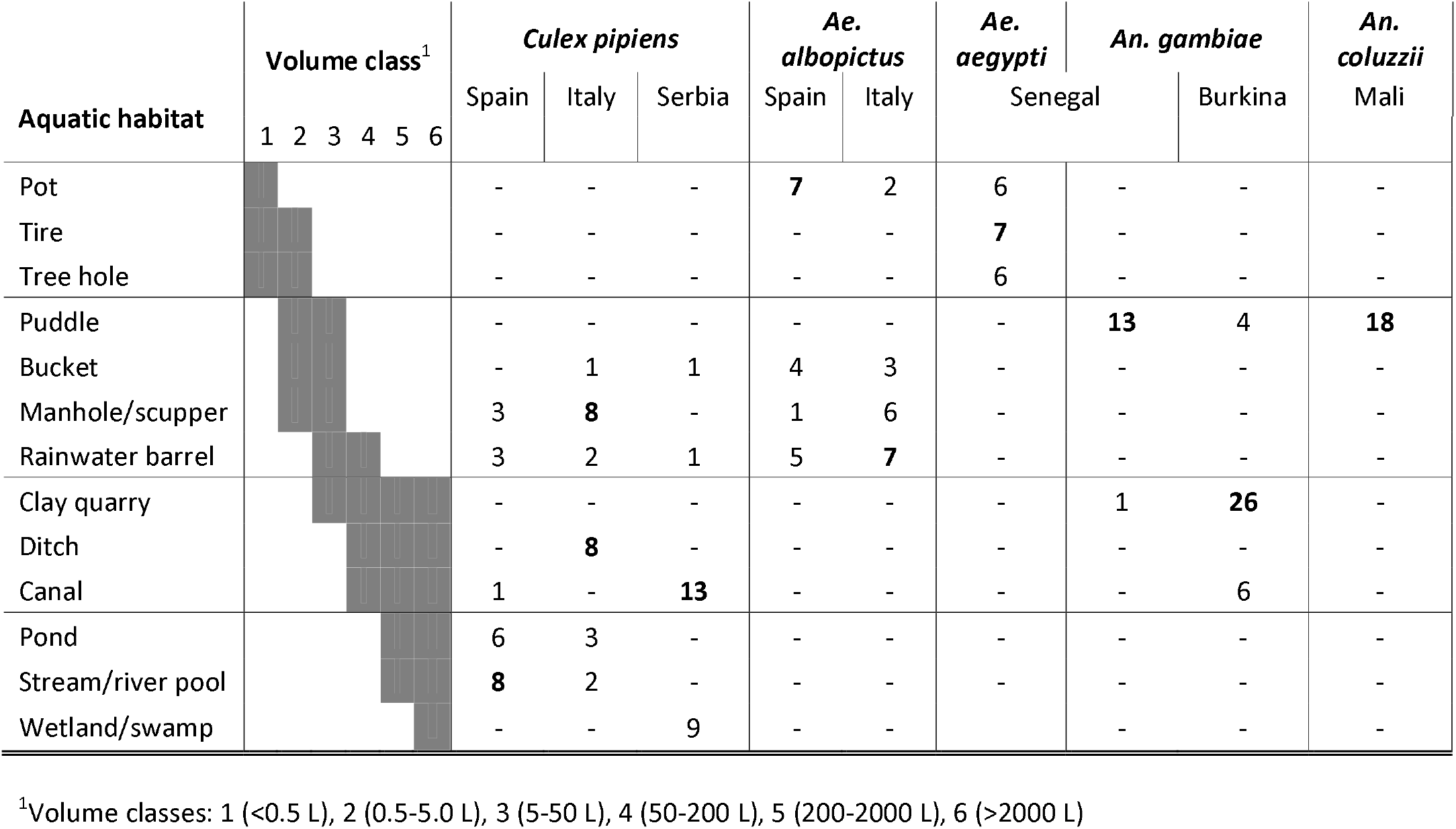
Typology of Mosquito breeding sites (MBS) of five target mosquito species including range of water volume (6 classes) as surveyed across six countries at two continents. The number of most common MBS types per species is marked in bold.

**Table 3.** Overall variation of water temperature [°C] was measured for the target species *Cx. pipiens*, *Ae. albopictus*, *Ae. aegypti*, *An. gambiae* and *An. coluzzii* in each mosquito breeding site. Next to the mean value per species, the overall variation is given as standard deviation (STD), coefficient of variation (CV), and minimum (Min) and maximum (Max) values for each species.

|  | *Cx.*<br>*pipiens* | *Ae.*<br>*albopictus* | *Ae.*<br>*aegypti* | *An.*<br>*gambiae* | *An.*<br>*coluzzii* |
| --- | --- | --- | --- | --- | --- |
| <u>Water temp. °C</u> |  |  |  |  |  |
| MEAN | 26.3 | 27.6 | 27.3 | 26.7 | 27.0 |
| STD | 4.1 | 3.7 | 1.6 | 2.6 | 2.2 |
| CV % | 15.7 | 13.3 | 5.8 | 9.9 | 8.0 |
| Min | 20.2 | 19.7 | 24.8 | 22 | 23 |
| Max | 37.6 | 37.1 | 31.3 | 31.8 | 30 |

**Fig. 2.**
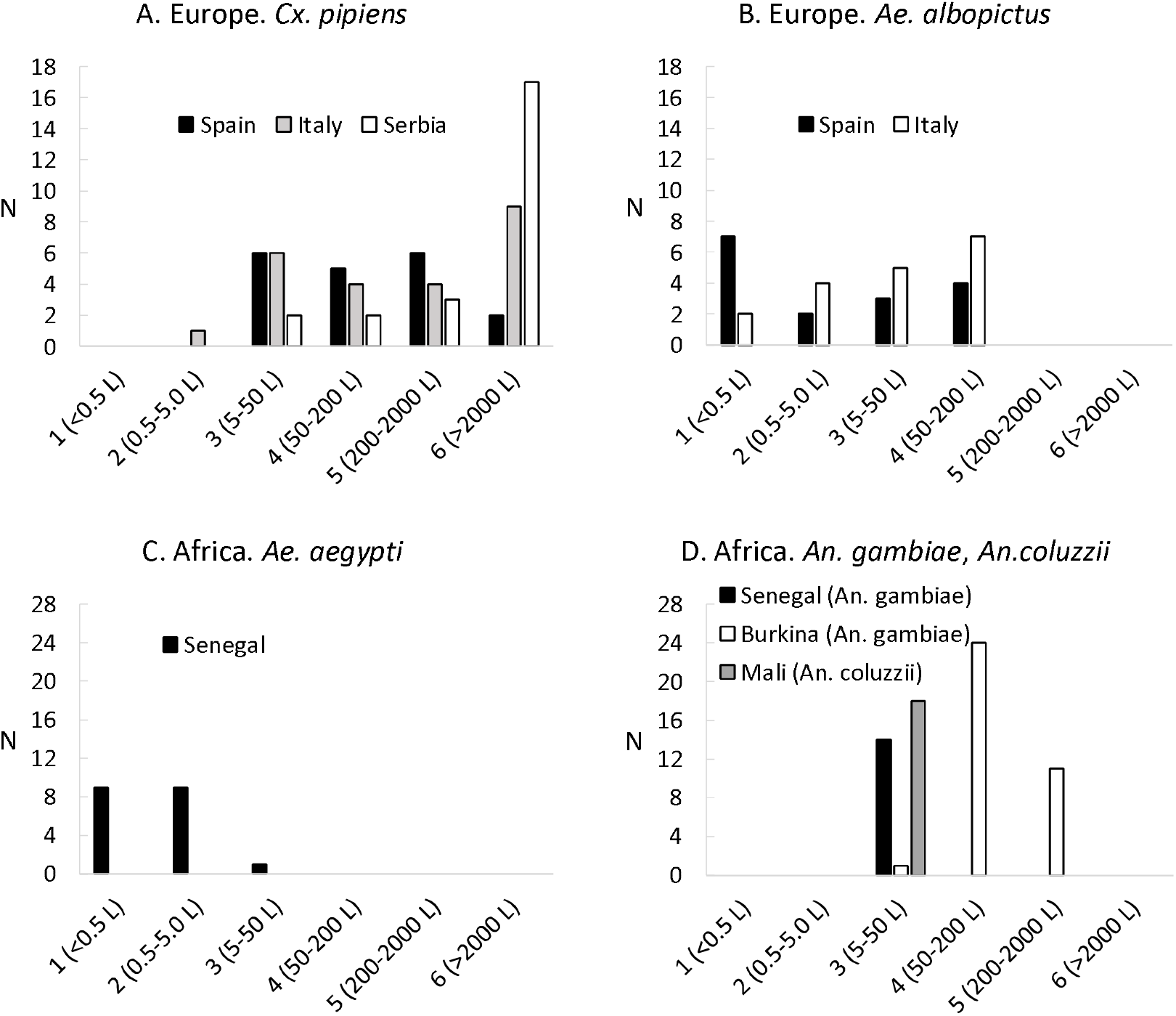
Number of aquatic individuals (larvae and pupae) per liter for five target mosquito species in six water volume classes of larval production sites as surveyed across six countries at two continents: *Cx. pipiens* (A), *Ae. albopictus* (B), *Ae. aegypti* (C), *An. gambiae* in Burkina Faso and Senegal and *An. coluzzii* in Mali (D).

**Fig. 3.**
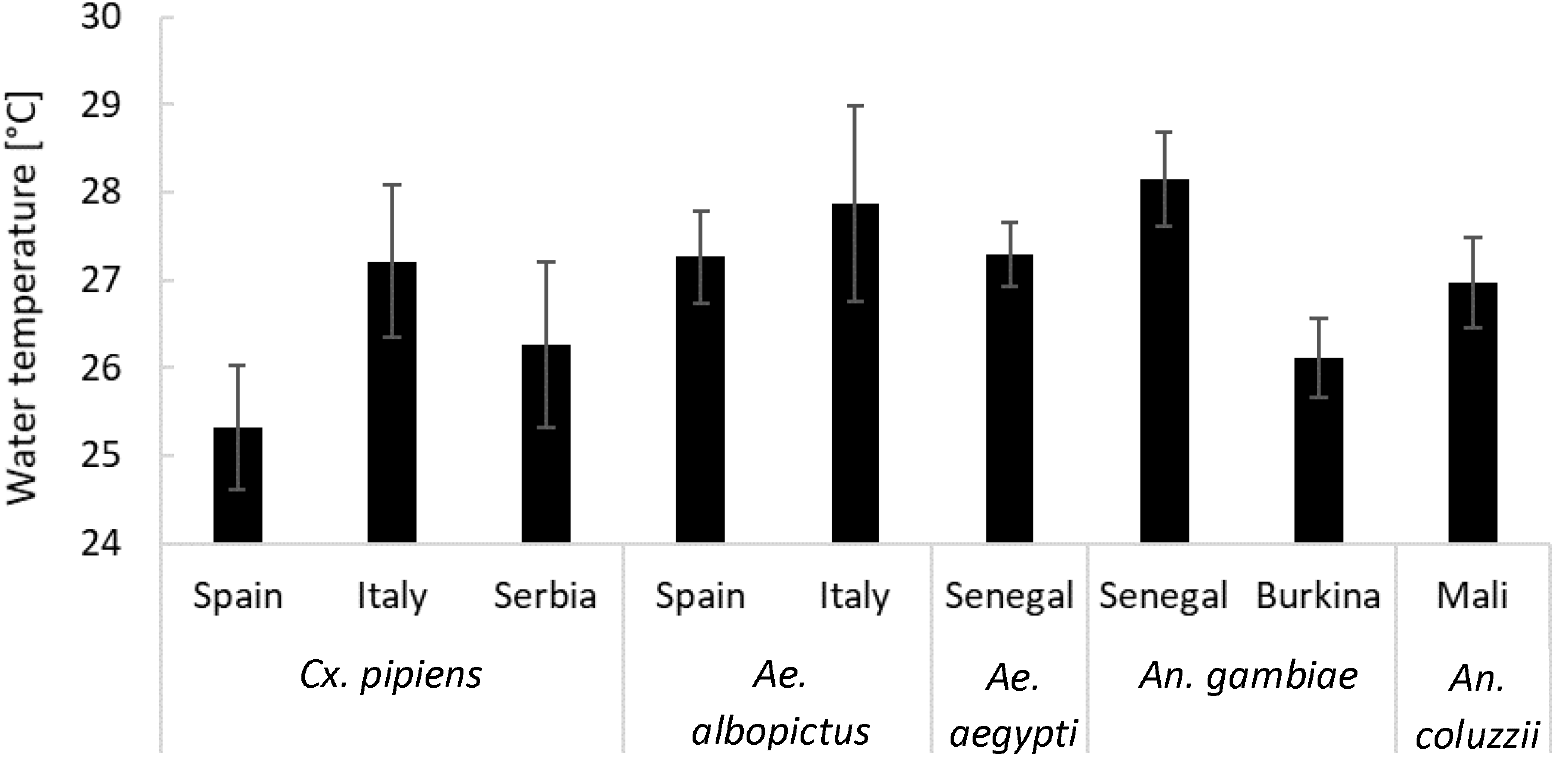
Water temperature [°C ± standard error] for five target mosquito species across six surveyed countries at two continents. Surveys were completed during peak season in European countries (July and August) and in Senegal and Burkina Faso (September to November), and at the end of the peak season in Mali (December).

The population density is highly variable across species though *Ae. albopictus* shows lower densities and smaller coefficient of variation compared to the other four taxa. The highest total population densities across countries are produced by *Cx. pipiens*. whereas its highest total density is reported from a waste container in Serbia (n=688 per liter). For comparison, the highest population density for *Ae. albopictus* was detected in a scupper drain in Spain (n=114 per liter). In Africa, *Ae aegypti* reaches 601 total individuals per liter in a tire in Senegal, *An. gambiae* 420 mosquitoes per liter in a puddle in Burkina Faso, and *An. coluzzii* 158 individuals per liter in a puddle in Mali.

The population structure described by larval and pupal density parameters shows a high coefficient of variation (CV) within each species (Table 4). The respective densities of L1-2 larvae, L3-4 larvae and pupae for each site is highly variable across countries (Fig. 4), though *Ae. albopictus* MBS in Italy and Spain show comparable proportions of pupae (18% and 20% respectively), and the proportions of late immature stages of *Cx. pipiens* MBS in Italy and Serbia are almost identical (12% for pupae and 42% for L3-4). The overall range of mean pupal abundance falls between 5.5 to 12.8 pupae per liter across all species (Table 4). Culex pipiens in Europe, *Ae. aegypti* and the two *Anopheles* species in Africa display very similar pupal ratios (0.10-0.12), while *Ae. albopictus* populations show a proportion of pupae as twice as high and the lowest variation (Table 4).

**Table 4.** The total number of aquatic life stages [mean (N larvae + pupae per liter)] and respective pupal ratio (N pupae / total N) per Mosquito breeding sites (MBS) were calculated for the target species *Cx. pipiens Ae. albopictus*, *Ae. aegypti*, *An. gambiae* and *An. coluzzii*. Together with the mean value per species, the overall variation among MBS is given as standard deviation (STD), coefficient of variation (CV), and minimum (Min) and maximum (Max) values for each species.

|  | *Cx.*<br>*pipiens* | *Ae.*<br>*albopictus* | *Ae.*<br>*aegypti* | *An.*<br>*gambiae* | *An.*<br>*coluzzii* |
| --- | --- | --- | --- | --- | --- |
| <u>Larval density (L1-2)</u> |  |  |  |  |  |
| MEAN | 71.8 | 19.1 | 41.4 | 55.1 | 38.6 |
| STD | 93.6 | 27.1 | 99.1 | 76.3 | 20.7 |
| CV % | 130.4 | 142.1 | 239.2 | 138.6 | 53.7 |
| Min | 0 | 0 | 0 | 0 | 10 |
| Max | 377 | 107 | 422 | 397 | 76.7 |
| <u>Larval density (L3-4)</u> |  |  |  |  |  |
| MEAN | 58.5 | 36.8 | 46.6 | 23.3 | 45.6 |
| STD | 92.0 | 9.8 | 101.4 | 22.9 | 22.8 |
| CV % | 157.2 | 26.6 | 217.4 | 98.4 | 50.1 |
| Min | 0 | 0.2 | 0 | 0 | 6 |
| Max | 475 | 118 | 430 | 96.0 | 85 |
| <u>Pupal density</u> |  |  |  |  |  |
| MEAN | 12.8 | 8.9 | 5.5 | 7.4 | 8.6 |
| STD | 27.7 | 9.8 | 8.8 | 5.5 | 10.6 |
| CV % | 216.4 | 109.6 | 160.5 | 74.1 | 122.2 |
| Min | 0.5 | 0.6 | 0 | 0 | 1 |
| Max | 204 | 33.5 | 26 | 25.0 | 40 |
| <u>Total population</u> |  |  |  |  |  |
| MEAN | 143.0 | 46.8 | 93.5 | 85.8 | 92.8 |
| STD | 180.1 | 40.4 | 175.4 | 91.8 | 42.4 |
| CV % | 125.9 | 86.3 | 187.6 | 106.9 | 45.7 |
| Min | 3 | 5 | 1 | 3 | 30 |
| Max | 928 | 123 | 601 | 420 | 157 |
| <u>Pupal ratio</u> |  |  |  |  |  |
| MEAN | 0.12 | 0.26 | 0.12 | 0.10 | 0.11 |
| STD | 0.14 | 0.20 | 0.26 | 0.09 | 0.14 |
| CV % | 117.1 | 79.0 | 211.0 | 86.5 | 123.7 |
| Min | 0.001 | 0.002 | 0 | 0 | 0.01 |
| Max | 1 | 0.90 | 0.29 | 0.33 | 0.55 |

**Fig. 4.**
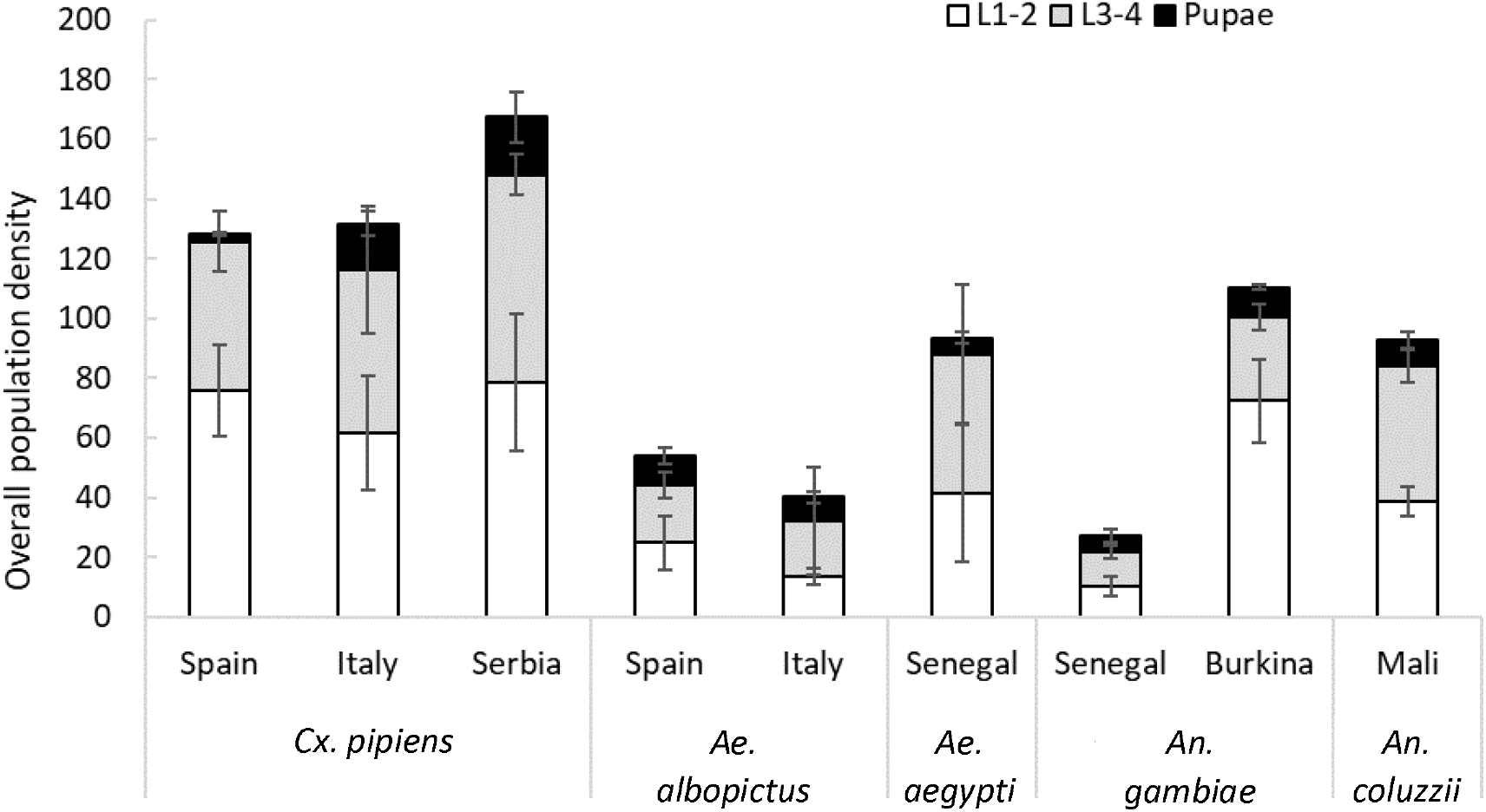
Larval (L1-2 and L3-4) and pupal densities per liter (mean ± standard error) for five target mosquito species surveyed in six countries at two continents. Surveys were completed during the peak season in European countries (July and August) and in Senegal and Burkina Faso (September to November), and at the end of the peak season in Mali (December).

## Discussion and conclusions

We successfully tested harmonized protocols to characterize the ecoregional variation of MBS characteristics and productivity for five mosquito vectors of primary public health importance in Europe and Africa. The protocols effectively capture variations in the preference for MBS and associated water volume, larval and pupal densities and water temperature with interesting patterns across species.

The overlapping presence of *Cx. pipiens* and *Ae. albopictus* in MBS with intermediate volume such as water barrels for irrigation and manholes highlights the widespread occurrence of larval competition in urbanized and rural environments (Carrieri et al. 2003). As both species infest small containers, niche partitioning for larval production and especially resource competition have been suggested (Carrieri et al. 2003, Marini et al. 2017, Müller et al 2018). Aedes aegypti preference for smaller sites (<5L), typically tires and tree holes, shows wide habitat availability for congeneric *Ae. albopictus* as it expands its invasive range in Africa (Simard et al. 2005). In Senegal and Mali we detected *An. gambiae* populations in large puddles, and in Burkina in pools formed in clay quarries. This confirms the ability of the two *Anopheles* species to colonize a variety of aquatic habitats, regardless of permanence (Edillo et al. 2002, Fillinger et al. 2004, Gimmonneau 2014). One may speculate that *An. gambiae* and *An. coluzzii* habitat preference seems purely driven by availability of water bodies rather than other site characteristics such as volume and temperature.

Interestingly, the range of mean water temperatures of MBS across species is small, between 26.3 and 27.6°C. This in vivo data on the range of large-scale temperature variation in MBS may improve the ability to design vector competence experiments in the laboratory, given that temperature during larval development is known to modulate vector competence (Westbrook et al. 2010, Murdock et al. 2014).

The variation of population densities and population structure is expectedly large within and across species inhabiting diverse MBS. The larval and pupal densities and their variation coincide with the density values reported for *Cx. pipiens* (Carrieri et al. 2013, Müller at al. 2018), *Ae. albopictus* (Eritja & Herreros, 2017), *Ae. aegypti* (Kamgang et al. 2013) and the two *Anopheles* species (Edillo et al. 2004, Sogoba et al. 2007). Interestingly, the mean pupal densities of five species range between 5 and 12 per liter and we assume, that those numbers could feed well mathematical models describing vector population dynamics. Also the mean pupal ratios of four out of five species (except *Ae. albopictus*) are commensurably: pupae constitute 10-12% of all life-stages with the exception of *Ae. albopictus* that shows a considerably higher pupal ratio, and the lowest variation. The unlike population structure and lower empirical variance might be related to the species adaptation as a colonizer of temporary waters. Optimal utilization of low resources leads to quicker larval development, which might skew *Ae. albopictus* population structure towards pupal stages (Marini et al. 2017), and lower predation pressure on immature stages in freshly colonized MBS could determine a higher overall MBS productivity.

As incidence of MBDs is globally rising, collaborative programs for insect vector surveillance at regional scale are crucial for predicting and managing outbreaks. Designing and implementing a multi-country mosquito vector monitoring program implies multiple organizational and ecoregional challenges. Limited manpower for professional field work in vector monitoring and control is a continuous challenge, thus reducing the temporal and geographical extent of areas monitored. In this study we demonstrate that collaborative projects among research groups are able to perform harmonized mosquito monitoring in two continents. We successfully employed harmonized protocols for characterizing mosquito MBS in six countries and over two continents. Thereby we confirmed that the available harmonized protocols represent a ready to use tool for implementing pan-regional vector surveillance programs and ecological studies. Our findings on population structures and densities might build a baseline for population modelling at continental scale for highly relevant mosquito vectors, and add important knowledge on species habitat preference and ecology that can be used in implementing IVM programs. Our results may also contribute to broaden the awareness of differences between the ways MBS are simulated in the laboratory against how they actually vary in the field.

## Acknowledgments

We thank Assétou Dionégué Diarra for data collection in Mali, Marta Verdun for technical support and data collection in Spain, and Dr Mihaela Kavran for technical assistance in Serbia, Marta Pagliochini for data entry, and Clause Meric for assistance with InfraVec web data download setup.

## Funding

This project has received funding from the European Union’s Horizon 2020 research and innovation programme Infravec2, Research infrastructures for the control of insect vector-borne diseases under grant agreement No 731060.

